# Genome-wide and high-density CRISPR-Cas9 screens identify point mutations in *PARP1* causing PARP inhibitor resistance

**DOI:** 10.1101/203224

**Authors:** Stephen J. Pettitt, Dragomir B. Krastev, Inger Brandsma, Amy Drean, Feifei Song, Radoslav Aleksandrov, Maria I. Harrell, Malini Menon, Rachel Brough, James Campbell, Jessica Frankum, Michael Ranes, Helen N. Pemberton, Rumana Rafiq, Kerry Fenwick, Amanda Swain, Sebastian Guettler, Jung-Min Lee, Elizabeth M. Swisher, Stoyno Stoynov, Kosuke Yusa, Alan Ashworth, Christopher J. Lord

## Abstract

PARP inhibitors (PARPi) target homologous recombination defective tumour cells via synthetic lethality. Genome-wide and high-density CRISPR-Cas9 “tag, mutate and enrich” mutagenesis screens identified single amino acid mutations in PARP1 that cause profound PARPi-resistance. These included PARP1 mutations outside of the DNA interacting regions of the protein, such as mutations in solvent exposed regions of the catalytic domain and clusters of mutations around points of contact between ZnF, WGR and HD domains. These mutations altered PARP1 trapping, as did a mutation found in a clinical case of PARPi resistance. These genetic studies reinforce the importance of trapped PARP1 as a key cytotoxic DNA lesion and suggest that interactions between non-DNA binding domains of PARP1 influence cytotoxicity. Finally, different mechanisms of PARPi resistance (*BRCA1* reversion, *PARP1*, *53BP1*, *REV7* mutation) had differing effects on chemotherapy sensitivity, suggesting that the underlying mechanism of PARPi resistance likely influences the success of subsequent therapies.

## Introduction

Drugs targeting the poly-(ADP-ribose) polymerase (PARP) enzymes PARP1 and PARP2 cause synthetic lethality in tumour cells with homologous recombination (HR) defects, including those with loss of function mutations in the *BRCA1* or *BRCA2* tumour suppressor genes^1–3^. PARP1 acts as a DNA damage sensor, rapidly binding single- and double-stranded DNA breaks as they occur and then coordinating their repair by synthesizing poly-(ADP-ribose) (PAR) chains on target proteins (PARylation)^4^. The rationale for using PARP inhibitors (PARPi) to treat HR-deficient cancers is based on the exquisite sensitivity of *BRCA1* or *BRCA2* defective cells to small molecule PARPi as well as the ability of *Parp1* gene silencing to selectively inhibit *Brca1* or *Brca2* defective cells^1,2^. Subsequent experiments have revealed that in addition to inhibiting the catalytic activity of PARP1, most clinical PARPi cause cytotoxicity by trapping PARP1 at sites of DNA damage^5–7^. The PARP1 trapping potency of different inhibitors correlates with their cytotoxic potency, with talazoparib (BMN673, Pfizer) showing the greatest effect^5,6^. Complete ablation of PARP1 expression by transposon-mediated mutagenesis or gene silencing in *BRCA1*/*BRCA2* wild type cells results in extreme resistance to several PARP inhibitors^1,7^.

The ability of some PARPi to trap PARP1 might be partially explained by the observation that PARP1 DNA binding is independent of its catalytic activity, while dissociation of PARP1 from DNA requires PARylation^8^. Recent structural studies have proposed a model of PARP1 binding to single stranded DNA damage that takes into account a series of molecular interactions between different PARP1 protein domains^9–11^. In its non-DNA bound state, a regulatory PARP1 helical domain (HD) is proposed to prevent catalytic activity. Upon PARP1 DNA binding (via N-terminal zinc finger DNA binding domains), an unfolding of the PARP1 helical region accompanies catalytic activation and PARP1 synthesizes PAR chains on itself and other acceptor proteins in the vicinity^9–11^. These PARylation events recruit other DNA repair enzymes, such as XRCC1^12^, and act as a second messenger signaling the presence of DNA damage. The synthesis of highly negatively-charged PAR chains on PARP1 is thought to also cause dissociation of PARP1 from DNA, presumably through a steric mechanism^8^.

Here we used CRISPR-Cas9 mutagenesis to investigate the mechanisms of PARPi toxicity in greater detail. We applied a focused mutagenesis approach to generate a large number of PARP1 mutant alleles that cause resistance, identifying an axis of intramolecular communication in PARP1 that mediates PARPi toxicity. We isolated PARP1 mutants from tumour cells with *BRCA1* exon 11 mutations and demonstrated that residual BRCA1 function in these cells allows tolerance of PARP1 loss of function despite the synthetic lethal relationship between these genes. A PARP1 mutation observed in a tumour from a PARPi resistant patient prevented PARP1 trapping, suggesting that *PARP1* mutations that impair trapping could contribute to clinical PARPi resistance. Finally, we found that *PARP1* mutations caused a distinct set of drug sensitivities when compared to other known forms of PARPi resistance (*REV7*, *53BP1* or *BRCA1* reversion mutants), suggesting that knowledge of the molecular mechanism of resistance in individual patients could inform decisions on further treatment.

## Results

### A genome-wide CRISPR screen for PARP inhibitor resistance identifies in-frame Parp1 mutants

Although PARPi are showing considerable promise as the first of a new generation of synthetic lethal therapies, resistance is a major issue^13,14^. To better understand this, we carried out a genome-wide CRISPR-Cas9 mutagenesis screen (encompassing 87,897 single guide (sg)RNAs) to identify mouse ES cell mutants resistant to the potent PARPi talazoparib^15^ (BMN673). We isolated and analysed 24 resistant clones (Methods, Figure 1a). Nine clones harboured one of two different sgRNAs targeting *Parp1* (Table 1). *Parp1* was the only gene that was targeted by more than one different sgRNA among the resistant clones (Table 1, Figure 1b).

**Figure 1.**
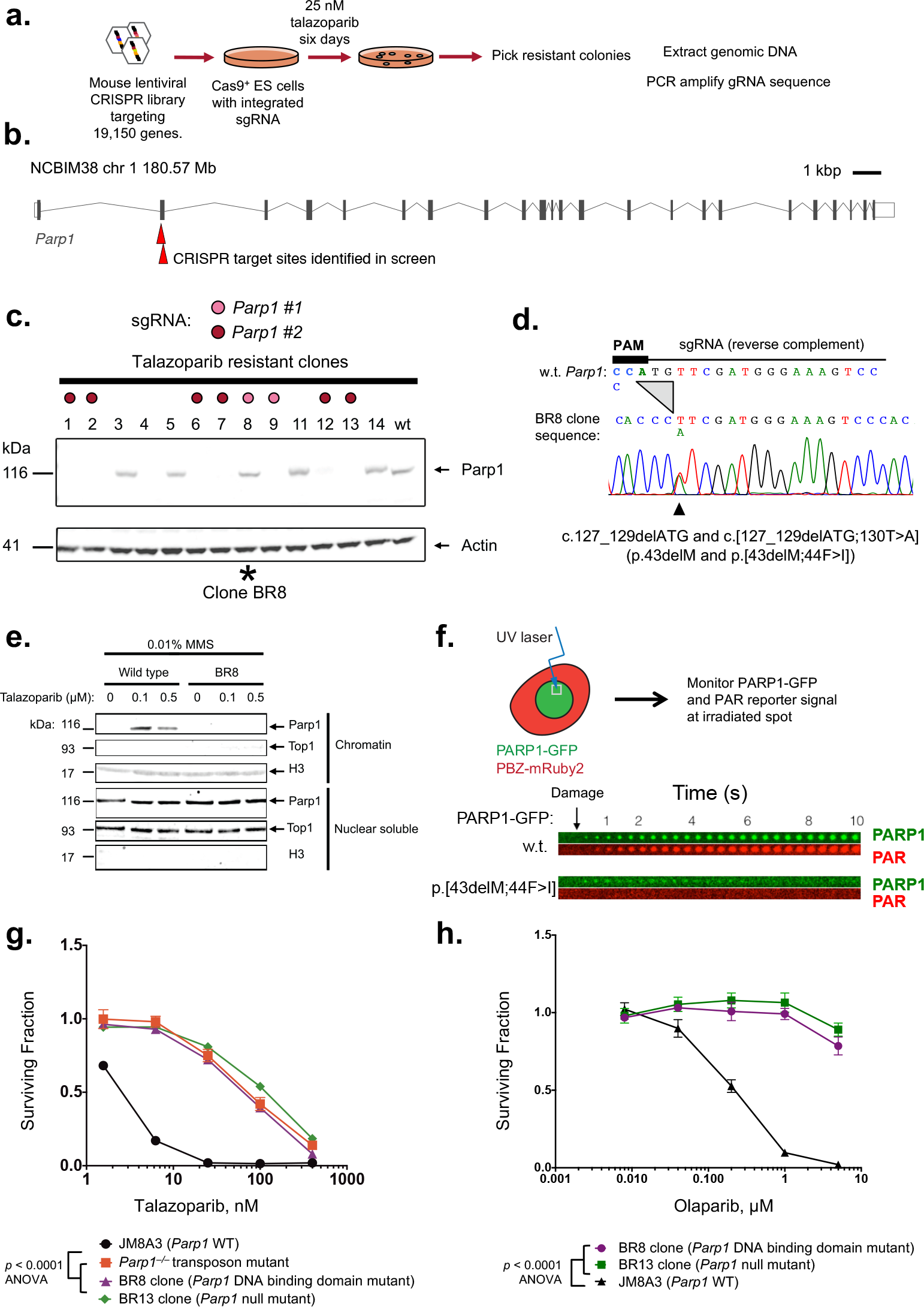
A genome-wide CRISPR screen for PARP inhibitor resistance identifies in-frame Parp1 mutants. **(a)** Experimental scheme: Cas9 expressing mouse embryonic stem (ES) cells were mutagenized with a CRISPR sgRNA library and PARP inhibitor resistant clones selected for by talazoparib exposure. sgRNA sequences in talazoparib-resistant clones were identified by PCR amplification/DNA sequencing. **(b)** Locations of *Parp1* guide RNA target sites in exon two of the mouse *Parp1* gene. **(c)** Both Parp1-expressing and Parp1 null clones are isolated from the genome-wide PARPi-resistance screen. Parp1 western blot of lysates from talazoparib-resistant clones identified in the CRISPR screen is shown. Individual clones are colour coded according to sgRNA present (see key). Clones 1, 2, 6, 7, 9, 12 and 13 with *Parp1* sgRNAs have lost Parp1 protein expression, whilst *Parp1* sgRNA clone 8 (BR8) has retained Parp1 expression. **(d)** The Parp1 expressing, talazoparib-resistant, BR8 clone has an in-frame Parp1 deletion and a Parp1 substitution mutation. Sanger sequencing trace from clone BR8 is shown, illustrating an in-frame Parp1 c.127_129delATG (p.43delM) homozygous deletion and a heterozygous c.130T>A substitution mutation (p.44F>I) close to the CRISPR PAM site. **(e)** Parp1 is not trapped in the chromatin fraction by PARP inhibitor in the BR8 clone. Western blots illustrating Parp1 in the chromatin and nuclear soluble fractions of wild type ES cells and Parp1 mutant BR8 cells exposed to talazoparib. Data shown are representative of two experiments. **(f)** PARP1 protein with a p.[43delM;44F>I] mutation has impaired recruitment to damaged DNA and does not initiate PAR synthesis at damaged DNA. Human CAL51 *PARP1*^−/−^; cells were transfected with either a wild-type PARP1-GFP or PARP1-p.[43delM;44F>I]-GFP fusion cDNA expression construct and then exposed to localised ionising radiation (a microirradiated spot). Temporal localisation of PARP1-GFP to damaged DNA was then estimated by visualising GFP signal at the microirradiated spot, as was the generation of PAR at the damaged site by use of a PAR binding PBZmRuby2 probe (see schematic). The time-course of PARP1-GFP and PBZmRuby2 signals from PARP1-GFP (top) and PARP1-p.[43delM;44F>I]-GFP transfected cells (bottom) are shown. “Damage” denotes time at which microirradiation was carried out. **(g)** and **(h)** The DNA binding domain BR8 clone is resistant to both talazoparib (g) and olaparib (h). Dose response curves illustrating PARP inhibitor resistance are shown. BR13 is a PARPi resistant Parp1 null ES cell mutant with no Parp1 protein expression (see panel b); a transposon-mutagenised, *Parp1* null mutant^7^ is also shown (red). Clone BR8 vs. BR13 and *Parp1*^−/−^ transposon, p = ns, ANOVA. Parp1 mutant clones vs. wild type cells, p < 0.0001 ANOVA. Mean of five replicates plotted, error bars show s.d.. Surviving fractions were calculated relative to DMSO exposed cells for each mutant.

**Table 1.**
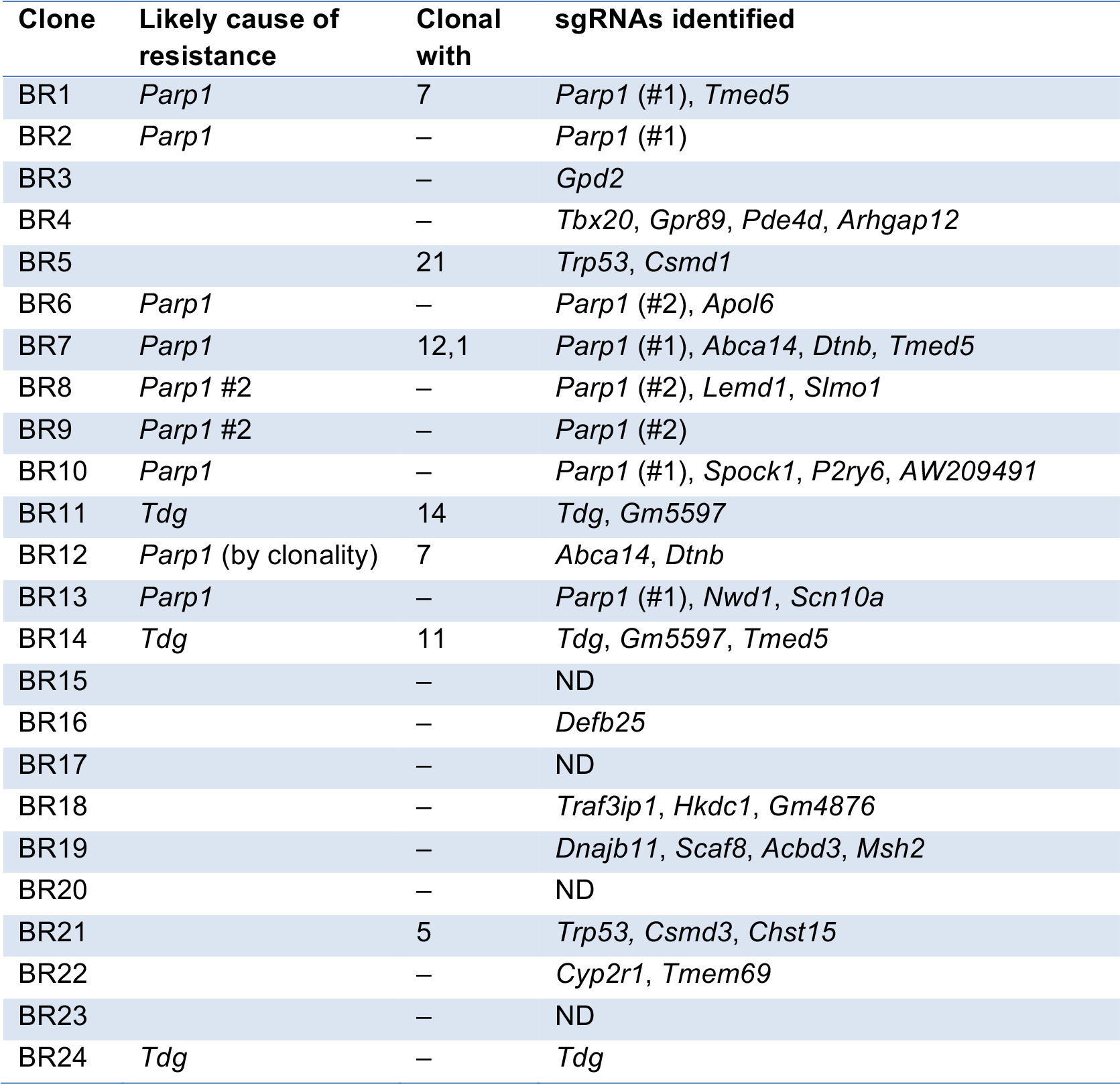
Results of sequencing sgRNA PCR products from talazoparib resistant mouse ES cell clones isolated from the CRISPR screen described in Figure 1a. Where clonality can be inferred from the combination of different guides observed, this is shown in column three. ND, not determined (failure of PCR and/or sequencing).

Parp1 protein was absent in all of the PARPi resistant clones with a *Parp1* sgRNA (Figure 1c) with one exception (clone BR8), consistent with the observation that ablation of PARP1 expression prevents PARP1 trapping and causes PARPi resistance^5,7^. DNA sequencing of the *Parp1* target site in clone BR8 revealed two in-frame *Parp1* mutations: c.127_129delATG and c[127_129delATG;130T>A] (p.43delM and p.[43delM;44F>I], Figure 1d). Both M43 and F44 residues are conserved between human and mouse PARP1 proteins and are predicted to be involved in base stacking interactions formed at the site of DNA/PARP1 interaction by the zinc fingers (ZnF) of the PARP1 DNA binding domain^16^. Using the PARP1 trapping assay^6^, we found that mutant Parp1 protein was not associated with the chromatin (C) fraction after talazoparib treatment, in contrast to the wild type protein (Figure 1e), suggesting that Parp1 trapping was impaired. A PARP1-GFP fusion protein with the p.[43delM;44F>I] mutation also failed to be recruited to DNA damage produced by laser microirradiation and did not produce poly-(ADP-ribose) (PAR) at the irradiated site, as monitored by expression of a fluorescent PAR binding reporter, PBZ-mRuby2 (Figure 1f). The magnitude of talazoparib resistance in the BR8 mutant clone was similar to that seen in *Parp1* mutants that showed complete loss of Parp1 protein expression, such as BR13, (Figure 1g). The BR8 clone also exhibited resistance to olaparib (Figure 1h, *p* <0.0001, ANOVA), suggesting a drug class effect. Taken together, these observations suggested that loss of Parp1 DNA binding and activity caused by mutations in the zinc finger domain of the protein can drive PARPi resistance.

### A focused CRISPR-Cas9 mutagenesis “tag, mutate and enrich” screen identifies PARP1 mutations outside the DNA binding domain that cause PARP inhibitor resistance

Our CRISPR screens highlighted a key feature of CRISPR mutagenesis – the ability to cause and easily identify subtle mutations as well as null mutations, such as the p.43delM mutation in the BR8 clone (Figure 1d). To study such mutations in more detail and to identify which regions of PARP1 are required for PARPi cytotoxicity, we designed an experimental approach to directly select PARPi-resistant cells with CRISPR-Cas9 induced mutations that preserved the native PARP1 reading frame (e.g. missense and in-frame indel mutations, rather than truncated, non-full length mutants). This approach, which we term, “tag, mutate and enrich” (Figure 2a), used HeLa cells with a BAC transgene containing *PARP1* gene fused to GFP coding sequence (encoding a PARP1 protein with a C-terminal GFP fusion). The GFP tag allowed us to enrich for cells with in-frame PARP1 protein expression by FACS isolating the GFP positive fraction. *PARP1-GFP*+ve cells were mutated with a focused sgRNA library of 29 guide RNAs targeting *PARP1* (Supplementary Table 1); PARPi resistant cells were then selected via talazoparib exposure and the GFP-positive fraction isolated (Figure 2a). The sgRNA library was introduced as six different lentiviral pools, grouped by RT-PCR product later used for genotyping. These products were amplified from GFP cDNA and sequenced using the Ion Torrent PGM platform (Figure 2a). For all but one sgRNA pool we saw a significant enrichment in in-frame mutations in *PARP1* in the PARPi resistant population (compared to a null hypothesis of 1/3 in-frame mutations; Pools 1, 2, 4, 5, 6: p < 10^−15^, binomial test with alternative hypothesis p(success) > 1/3, 95% CI 0.62-1.00. Pool 3, 95% CI 0.24-1.00, p = 1; Supplementary Figure 1a). By translating and aligning in-frame reads from PGM sequencing, we identified key PARP1 amino acid residues associated with resistance. For example, we identified multiple alleles in the PARPi resistant population with deletion of nucleotides encoding amino acid residues K119 and S120 (Figure 2b), which are both DNA-contacting residues within the second zinc finger domain of PARP1 (reference^16^). Our subsequent microirradiation assay analysis demonstrated that a p.119_120delKS mutation abolished PARP1-GFP recruitment to sites of DNA damage (Figure 2c), suggesting that mutations in residues K119 and S120 might cause PARPi resistance by altering PARP1 DNA binding properties.

**Figure 2.**
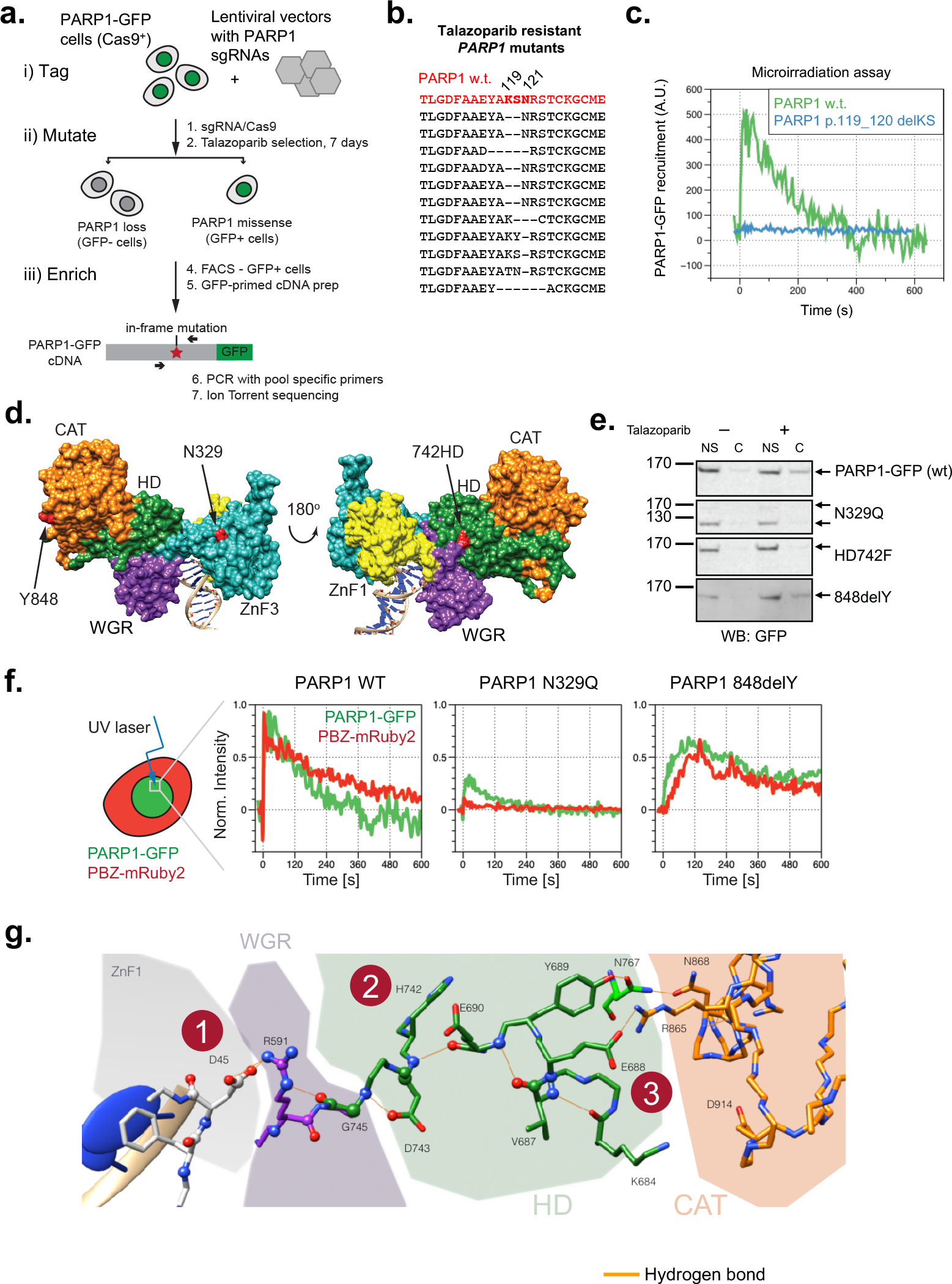
A focused CRISPR-Cas9 mutagenesis “tag, mutate and enrich” screen identifies PARP1 mutations outside the DNA binding domain that cause PARP inhibitor resistance. **(a)** Experimental “tag, mutate and enrich” scheme to isolate missense and in-frame *PARP1* mutants associated with PARP inhibitor resistance. A Cas9-expressing HeLa clone with a transgenic bacterial artificial chromosome containing the *PARP1* gene, fused at the 3′ terminus to a GFP coding sequence (PARP1-GFP “tagged” cells) were infected with lentiviral pools of *PARP1*-targeting lentiviral sgRNA expression constructs (“mutate”). In total, six different lentiviral pools were used; each sgRNA pool targeted a different region of *PARP1* as shown in (a). *PARP1* mutant cells were selected from this population by exposing cells to talazoparib. To enrich for *PARP1* mutant clones likely to have either missense or in-frame PARP1 indel mutations (and not premature truncating mutations), PARP1-GFP positive cells were selected by FACS. *PARP1* mutations in the GFP-positive, talazoparib-resistant cell populations were identified by RT-PCR and IonTorrent sequencing. **(b)** Translated alignment of PARP1 amino acid mutations identified in talazoparib resistant cells isolated from lentiviral sgRNA Pool 1 (designed to target ZnF1 and ZnF2 domains) from the screen shown in A. By comparing the multiple different mutations isolated from Pool 1, a PARP1 p.119_120delKS minimal mutation associated with resistance was identified. Some insertions and larger deletions are omitted for clarity (see Supplementary Table 2). **(c)** PARP1 p.119_120delKS mutation abolishes recruitment of a PARP1-GFP fusion to sites of microirradiated DNA damage. CAL51 *PARP1*^−/−^ cells were transfected with a wild-type PARP1-GFP cDNA construct or a PARP1 p.119_120delKS-GFP fusion cDNA expression construct and then exposed to localised ionising radiation (a microirradiated spot) as in Figure 2f. Time course of PARP1-GFP signal at microirradiated site is shown. **(d)** Location of three mutations associated with PARPi resistance on a model of the PARP1-DNA structure^10^. **(e)** PARP1 N329Q and HD742F mutations ablate PARP1 trapping, while 848delY partially reduces trapping. Western blot from PARP1 trapping assay for three PARP1-GFP mutants and wild type PARP1-GFP is shown. MMS-treated cells were lysed and fractionated into nuclear soluble (NS) and chromatin (C) fractions as described in Methods. Blot was probed with an anti-GFP antibody. **(f)** PARP1 848delY mutation alters PARP1 localisation kinetics at sites of DNA damage. Microirradiation and PAR synthesis phenotypes of N329Q and 848delY PARP1 mutants. Green – PARP1-GFP signal, red – PAR sensor (PBZ-mRuby2). Note lower recruitment relative to wild type of both mutants, but retention of PAR synthesis in 848delY. **(g)** Model of intramolecular communication between the DNA binding ZnF1 domain and the catalytic (CAT) domain based on mutants identified from screens. (1) 45delD mutation in first zinc finger (2) HD742F mutation described in (e), (3) E688.

The focused sgRNA screen also identified three *PARP1* mutations in residues not known to be directly involved in DNA binding: p.329N>Q (N329Q), p.742-743HD>F (HD742F) and p.848delY (848delY, Figure 2d, Supplementary Tables 2 and 3). N329 sits within the third zinc finger of PARP1, but is not predicted to form part of the DNA binding interface. HD742F is located within the helical domain (HD), a regulatory region shown to be important for PARP activation^9,10^. Y848 is part of a solvent-exposed helix of the catalytic domain (Figure 2d). In cell-free assays using recombinant PARP1 proteins, we found the talazoparib IC_50_ to be similar for mutant and wild type PARP1 proteins (Supplementary Figure 1b, c), suggesting that the PARPi resistance phenotypes were not caused by differences in the ability of PARPi to inhibit the catalytic activity of mutant proteins. PARP1 trapping assays in CAL51-*PARP1*^−/−^ cells transfected with mutant or wild type (WT) PARP1-GFP cDNA expression constructs suggested that N329Q and HD742F mutant proteins were not trapped by PARPi (Figure 2e), explaining the PARPi resistant phenotypes associated with these mutations. In contrast, PARP1-848delY-GFP was trapped in the chromatin fraction by talazoparib, but to a lesser extent than WT-PARP1-GFP (Figure 2e). Laser microirradiation assays confirmed this partial PARP1 trapping phenotype with the PARP1-848delY-GFP protein; whilst WT-PARP1-GFP was rapidly recruited to the site of microirradiation (peak ≈ 4s) and induced PAR formation in a similar time frame (Figure 2f), the extent of PARP1-848delY-GFP recruitment to the site of microirradiation was reduced, as was PARylation (Figure 2f). As a negative control, PARP1-N329Q-GFP exhibited limited recruitment to damaged DNA (Figure 2f). The addition of talazoparib delayed PARP1-848delY-GFP dissociation from microirradiated sites, but to a lesser extent than for WT-PARP1-GFP (Supplementary Figure 1d, p = 5 × 10^−7^, t-test). PARP1-848delY-GFP also dissociated faster from microirradiated sites, being absent from microirradiated regions 30 minutes after microirradiation; WT PARP1-GFP still showed 40% of maximal trapping at this timepoint (Supplementary Figure 1e, p = 5.5 × 10^−3^, t-test). These data therefore suggested that even the relatively subtle defect in trapping in the 848delY mutant (Figure 2e, f) might be sufficient to cause PARPi resistance.

We mapped the residues affected by the mutations that we observed onto a previously described crystal structure of PARP1 zinc finger domains 1 and 2 together with the WGR, regulatory and catalytic domains^11^. We found that several of the PARPi resistance-causing mutations (D45, H742, D743, E688) affected amino acid residues involved in hydrogen bonding interactions that bridge the DNA binding domain and the catalytic domain (Figure 2g), suggesting that these might control inter-domain interactions that mediate PARP1 trapping; similar inter-domain domain interactions link DNA binding to activation of PARP1 catalytic activity^9–11^. Taken together, these data established that mutations outside of the DNA binding domain can cause PARPi resistance, likely by impairing PARP1 trapping.

### *PARP1* mutations cause PARPi resistance in *BRCA1* mutant tumour cells with mutations in exon 11

We assessed whether *PARP1* mutations could cause PARPi resistance in a clinically relevant setting, such as in *BRCA1* mutant tumour cells. Complete loss of both *PARP1* and *BRCA1* is expected to be synthetic lethal^1,2^. However, a growing body of evidence suggests that many pathogenic *BRCA1* mutations may not result in complete loss of function. For example, SUM149 (also known as SUM149PT) triple negative breast tumour cells possess a commonly-occurring hypomorphic *BRCA1* exon 11 c.2288delT frameshift mutation and loss of the wild type *BRCA1* allele^17^. As a consequence, SUM149 cells do not express the full length BRCA1 p220 protein, but do express a hypomorphic splice isoform of BRCA1, Δ11b, which excludes the c.2288delT premature truncating mutation (along with most of the exon 11 coding sequence), but has some residual function^18,19^. We and others have previously confirmed that the *BRCA1* mutation in SUM149 cells causes sensitivity to PARPi by demonstrating that genetic reversion of the *BRCA1* mutation in SUM149 cells via CRISPR-Cas9 mutagenesis imparts PARPi resistance^19,20^.

We carried out a genome wide PARPi resistance CRISPR-Cas9 screen in SUM149 cells, using a previously validated sgRNA library^21^, similar to our earlier screen in mouse ES cells. Out of 12 talazoparib-resistant clones analysed, eight possessed one of three PARP1 sgRNAs (Table 2), suggesting that talazoparib cytotoxicity is mediated by PARP1 even in this *BRCA1* mutant cell line. To confirm this observation, we infected Cas9-expressing SUM149 cells with a lentiviral sgRNA vector targeting the *PARP1* coding sequence homologous to the p.43/44 mutation site in the PARPi resistant mouse ES cell BR8 clone we identified earlier (Figure 1d). This sgRNA induced talazoparib resistance (Figure 3a). We subcloned two daughter clones from the PARPi-resistant SUM149 population, TR1 and TR2; TR1 had four different *PARP1* frameshift mutations and lacked PARP1 protein expression, whilst TR2 expressed apparently full length PARP1 protein (Figure 3b). Both clones remained talazoparib-resistant after culture in the absence of PARPi, demonstrating that this was a stably-acquired phenotype (Figure 3c, ANOVA p < 0.0001). TR2 possessed three different frameshift PARP1 mutations (Table 3) but also a p.43_45delMFD in-frame deletion, similar to the p.43/44 mutations identified in PARPi resistant BR8 mouse ES cells (Figure 1d). No *PARP1* wild type DNA sequencing reads were identified in either TR1 or TR2. The allele frequencies of the *PARP1* mutations in TR1 and TR2 suggested that there were five copies of the *PARP1* locus in SUM149 cells, confirmed by SNP genotyping of SUM149 (A. Grigoriadis, personal communication). We did not find evidence for other known mechanisms of PARPi resistance in TR1 and TR2 such as reversion of full-length BRCA1 expression^22^ (Supplementary Figure 2a), 53BP1 or REV7 (MAD2L2) loss^23^ (Supplementary Figure 2b, c), or an increase in DNA damage-induced nuclear RAD51 foci (Supplementary Figure 2d). This suggested that PARP1 mutation or loss could not only be tolerated in SUM149 cells but also caused PARPi resistance.

**Table 2.** sgRNA sequences identified in talazoparib-resistant SUM149-Cas9 cells mutagenized with a genome-wide CRISPR library.

| sgRNA | Colonies |
| --- | --- |
| PARP1_ex16 | 5 |
| PARP1_ex19 | 1 |
| PARP1_ex22 | 1 |
| PARP1_ex19, MCF2L_ex2 | 1 |
| PARP1_ex22, JPH4_ex3 | 1 |
| RABGAP1_ex8 | 1 |
| ND (>2 guides) | 2 |
|  | 12 |

**Figure 3.**
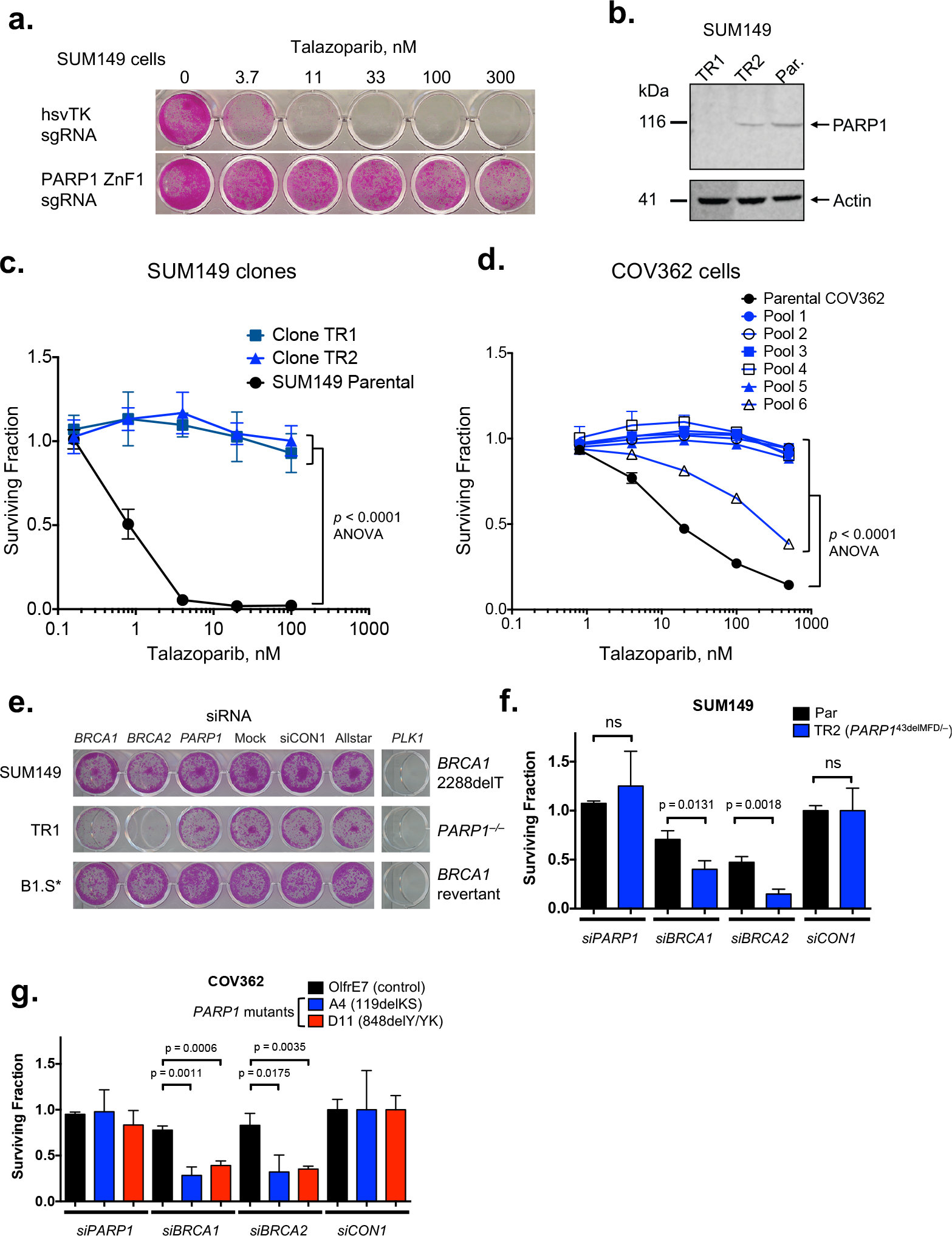
*PARP1* mediates PARPi sensitivity in *BRCA1* mutant cell lines. PARP1 ZnF1 guide lentiviral infection drives talazoparib resistance in *BRCA1* mutant SUM149 triple negative breast cancer cells. Cas9-expressing cells were transduced with lentiviral sgRNA particles, selected in puromycin and exposed to the indicated talazoparib concentrations for seven days. Representative crystal violet stained 24 well plate image after is shown. An sgRNA lentivirus targeting hsvTK (herpes simplex virus thymidine kinase, not present in the human genome) was used as a negative control. Numbers above each well indicate the concentration of talazoparib (nM). **(b)** Western blot showing PARP1 expression in talazoparib resistant SUM149 clones TR1 and TR2. Clone TR1 lacks detectable PARP1. Par, parental SUM149 cells. **(c)** SUM149 clones TR1 and TR2 are highly talazoparib resistant. Plot shows survival relative to DMSO exposed cells after exposure to the indicated concentration of talazoparib for seven days. Compared to the response in the parental clone, both clone TR1 and TR2 are significantly resistant (p<0.0001 ANOVA). Mean of five replicates plotted, error bars show s.d.. **(d)** Pools of *PARP1* guides cause talazoparib resistance in the *BRCA1* mutant ovarian cancer cell line COV362 (*BRCA1* c.2611fs/c.4095+1G>T). Cas9-expressing COV362 cells were transduced with the indicated lentiviral guide pool, selected in puromycin and exposed to the indicated talazoparib concentrations for seven days. ANOVA *p* < 0.0001 compared to parental cells for all pools. Mean of five replicates plotted, error bars show s.d.. Surviving fractions were calculated relative to DMSO exposed cells for each mutant. **(e)** Silencing of residual *BRCA1* function, or *BRCA2*, induces synthetic lethality with *PARP1* genetic loss in SUM149 cells. B1.S* is a SUM149 derivative with a secondary *BRCA1* mutation that confers PARP inhibitor resistance. Crystal violet stained 24 well plate image is shown, seven days after siRNA transfection. **(f)** Colony forming assay using SUM149-Cas9 parental cells and the talazoparib resistant daughter clone, TR2. Cells were transfected as in (f), plated on six-well plates and colonies stained and counted two weeks later. siRNA targeting *BRCA1* or *BRCA2* significantly reduces survival in *PARP1* mutant cells (TR2) compared to the parental SUM149 line (t-test). Mean of three replicates plotted, error bars show s.d.. **(g)** Colony forming assay for COV362 *PARP1* mutants. *BRCA1* and *BRCA2* silencing is also synthetically lethal in COV362 *PARP1* mutant clones A4 (p.119_120delKS) and D1 (p.848delY/YK). Colony formation assay as in (g), p values for t-test shown. Mean of three replicates plotted, error bars show s.d..

**Table 3.** *PARP1* mutations identified in talazoparib-resistant SUM149 and COV362 clones, determined by Sanger sequencing. *, stop codon; fs, frame shift resulting in the indicated number of different amino acid residues.

| Clone | sgRNA | *PARP1* mutation(s) |
| --- | --- | --- |
| SUM149 TR1 | ZnF 1 | Frameshift mutations: c.130_133delTTTG (p.[44delF;fs49*]), c.130_131delTT (p.44F>*), c.129_130delGT (p.43M>I*), c.121-4_130_delCTAGTCGCCCATGTT [splice acceptor] |
| SUM149 TR2 | ZnF 1 | p.43delMFD and frameshift mutations: c.130_133delTTTG (p.[44delF;fs49*]), c.129_130insC (p.44F>L*), c.129delG (p.43M>fs49*). |
| COV362 A4 | Pool 2 | p.119delKS |
| COV362 D1 | Pool 5 | p.848delY/848delYK |
| COV362 D4 | Pool 5 | p.848delYKPF |
| COV362 D8 | Pool 5 | Wild type |
| COV362 D9 | Pool 5 | p.848delY/848delYK |
| COV362 D11 | Pool 5 | p.848delY/848delYK |

We went on to show that *PARP1* sgRNA also caused PARPi resistance in *BRCA1* mutant COV362 ovarian tumour cells^24^ (*BRCA1* c.2611fs [exon 11] and c.4095+1G>T [exon 11 splice donor]), suggesting that these observations were not private to SUM149 cells. COV362 cells were infected with six different pools of lentiviral *PARP1* sgRNAs as used in the HeLa screen above (Figure 2) and selected in talazoparib. We found profound talazoparib resistance in all lentivirally-infected sgRNA-infected populations, except those infected with *PARP1* sgRNA pool six, where resistance was less pronounced, although still significant (Figure 3d; p < 0.0001 in each case, ANOVA compared to parental COV362-Cas9 cells). We also isolated a number of daughter clones from the PARPi resistant COV362 populations. Some of these clones had lost PARP1 expression while others had point mutations in PARP1, including a clone with a p.119delKS mutation and three independently derived clones all with p.848delY/p.848_849delYK compound mutations (referred to as p.848delY/YK, Supplementary Figure 2e, Table 3). All clones with *PARP1* mutations remained resistant to talazoparib after culture without PARPi selection during subcloning and expansion (Supplementary Figure 2f, g). Interestingly, COV362 clones isolated with Y848 mutations at the endogenous *PARP1* locus (p.848delY/delYK) showed a less pronounced resistance phenotype compared to complete null clones, although the extent of resistance was still significant compared to wild type cells (Supplementary Figure 2f, g, h). These clones also showed some residual PARylation activity in agreement with the microirradiation phenotype observed earlier (Supplementary Figure 2i, Figure 2f), suggesting an intermediate level of PARPi resistance could be caused by an partial PARP1 trapping defect.

### Residual *BRCA1* function supports *PARP1* mutations in tumour cells

We also attempted to isolate *PARP1* mutants from MDA-MB-436 cells, which have a *BRCA1* c.5396 + 1G>A mutation in the splice donor site of exon 20. This mutation results in a truncated BRCA1 protein lacking the BRCT region required for HR^17,25^. We were unable to isolate long-term talazoparib resistant cells from this cell line after transfection with PARP1 sgRNA expressing lentiviral pools. Analysis of cells grown in talazoparib for a short period showed that mutations were induced at the sgRNA target sites, but cells with PARP1 mutations did not survive in the long term (Supplementary Figure 2j, k). In contrast, sequencing of COV362 cells infected with the same PARP1 sgRNA pools revealed that several pools of talazoparib-resistant cells generated in the same way had mutagenized all *PARP1* alleles in all cells, indicated by very low or absent wild type reads (Supplementary Figure 2k). This suggested that mutation of all *PARP1* alleles was not tolerated in MDA-MB-436 cells, in contrast to SUM149 or COV362 cells.

We therefore considered it possible that the residual *BRCA1* function in SUM149 and COV362 might allow these cells to tolerate *PARP1* mutations. To address this, we inhibited the residual BRCA1 function in SUM149 and COV362 cells with siRNA targeting *BRCA1* target sites outside exon 11; doing the same in SUM149/COV362 *PARP1* mutants isolated earlier demonstrated that *BRCA1* siRNA had a more profound cell inhibitory effect in *PARP1* mutant COV362 and SUM149 cells, than in *PARP1* wild type parental cells (Figure 3e-g, p < 0.05 for all cases compared to parental cells, t-test). This suggested that some residual BRCA1 function might exist in these cells and is required for cell survival in the face of *PARP1* mutation. *BRCA2* siRNA also selectively targeted *PARP1* mutant COV362 and SUM149 cells, suggesting some requirement for BRCA2 function in these cells despite the *BRCA1* mutation (Figure 3e-g, p < 0.05, t-test). Taken together, these observations suggested that *PARP1* loss might be tolerated in COV362 and SUM149 cells due to some residual *BRCA1*-dependent function. Exon 11 frameshift mutations comprise approximately 30% of pathogenic *BRCA1* mutations identified to date^19,26–29^ and thus *PARP1* mutation and/or loss could be a clinically relevant mechanism of PARPi resistance in this specific context.

### A high-density tiling CRISPR-Cas9 screen enables functional annotation of a *PARP1* mutation observed in an olaparib-resistant patient

To investigate which functions of PARP1 were important for cytotoxicity of talazoparib in *BRCA1* mutant cells, we designed a new tiling CRISPR library, comprising 489 guides, designed to give the densest possible coverage of PARP1 mutations (Supplementary Table 4, Figure 4a) and carried out a tiling screen in SUM149 cells in which the endogenous PARP1 locus had been tagged with a C-terminal GFP coding sequence using the CRISPaint system^30^. This screen further illuminated the roles of different areas of PARP1 in PARPi toxicity. In agreement with our previous results, a large number of mutations were isolated which affected residues associated with either direct DNA binding (Figure 4b, c; Supplementary Table 5) or the interfaces between zinc finger domains (Figure 4d). We also observed a high density of mutations affecting the WGR domain associated with PARPi resistance (Figure 4b, e). Based on our previous screen, we hypothesized that points of contact of the WGR domain with the zinc finger and HD (regulatory) domains might be important for PARPi-resistance and PARP1 function and activation (Figure 2f); in this new screen we identified several mutation clusters at these points of contact (Figure 4b, e; Supplementary Figure 3a) supporting this hypothesis. Although we identified few mutations in the catalytic domain that caused PARPi resistance – perhaps because mutations directly disrupting the catalytic activity would cause constitutive trapping and cytotoxicity – the most frequent mutation isolated in the screen affected residue A925, which is juxtaposed with residue Y848 in the three-dimensional structure of PARP1, further suggesting an important role for this region of the catalytic domain in PARPi cytotoxicity (Figure 4e, inset 2). The relative lack of mutations in the BRCT domain, despite ample sequencing and sgRNA coverage in this area (Figure 4c and Supplementary Figure 3b) might indicate that this region is dispensable for PARPi cytotoxicity and trapping.

**Figure 4.**
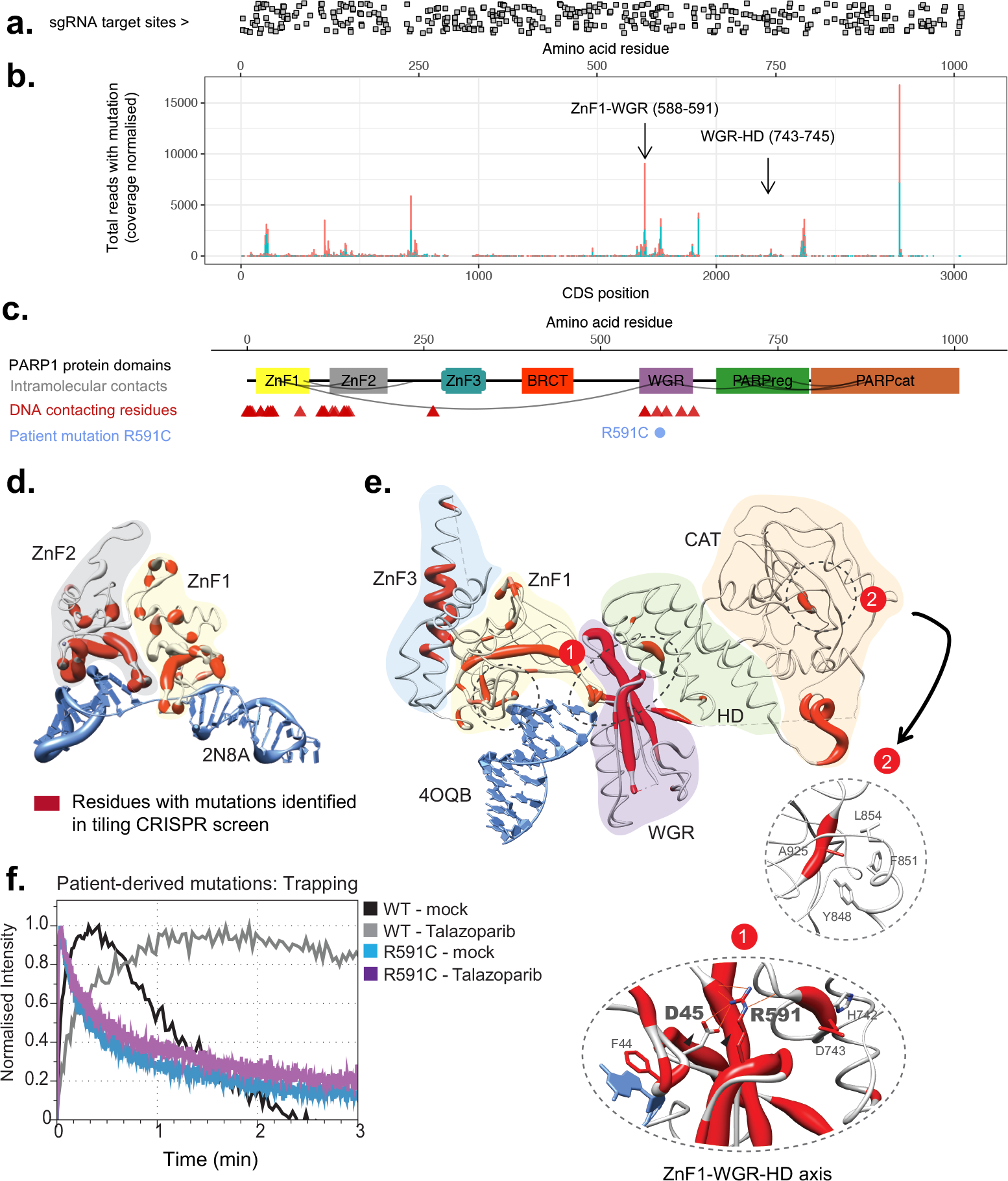
A high-density tiling CRISPR-Cas9 screen enables functional annotation of a *PARP1* mutation observed in an olaparib-resistant patient. **(a)** Positions of sgRNA target sites in the dense PARP1 tiling library mapped onto the PARP1 coding DNA sequence shown below in (b). **(b)** Positions of mutations identified in the dense *PARP1* tiling screen. Total number of mutant reads for each in-frame mutant allele affecting each base is shown on the y-axis (log scale), normalised to per-base coverage. The experiment was repeated in duplicate from two independently-tagged SUM149 PARP1-GFP cell lines (clone 5 and 8, shown in red and blue). Arrows highlight the mutations observed at interdomain contacts as shown in (e) and (g), below. **(c)** Positions of protein domains mapped onto the PARP1 protein sequence. Red triangles indicate DNA-protein contacts based oncrystal structures, blue circles show the location of the patient mutation identified in this study. Grey arcs represent key interdomain contacts as shown in (g) and Figure 2g. **(d)** Ribbon plot of ZnF1 and ZnF2 bound to a single strand break (PDB: 2N8A). Ribbons are coloured red by residue based on the frequency of mutations observed in the tiling screen affecting that residue as shown in (b). The thickness of the ribbon is proportional to the frequency of mutations observed affecting that residue. **(e)** Ribbon plot of the PARP1-DSB crystal structure (PDB:4OQB) highlighting the clustering of mutations along the hydrogen bonding axis postulated in Figure 2g and the A925 residue that abuts Y848 (inset 2). Regions marked 1 and 2 are magnified. **(f)** Analysis of trapping for the R591C patient mutation by microirradiation with or without talazoparib as shown. Although the R591C mutant can bind DNA at sites of damage (blue line), it is not trapped in the presence of talazoparib (purple line).

In parallel with these genetic studies, we identified *a PARP1* p.R591C mutation (c.1771C>T) in an ovarian cancer patient who showed *de novo* resistance to olaparib. The R591C mutation affects the WGR domain at the point of contact with D45 and the HD domain; our model (Figure 2g) and previous work^9–11^ predicts that this would be a critical residue for inter-domain communication in PARP1, between the DNA binding and catalytic domains, implying that this mutant might bind DNA but would not show trapping. To assess this, we assessed the recruitment of a PARP1-R591C-GFP fusion protein to sites of micorirradiation in the presence and absence of the PARPi talazoparib. We found that whilst PARP1-R591C-GFP was recruited to sites of microirradiation, its dissociation from these sites was rapid compared to wild type PARP1-GFP; furthermore, PARP1-R591C-GFP was not trapped on DNA by talazoparib, unlike wild type PARP1-GFP (Figure 4f).

### Distinct vulnerabilities of cells acquiring PARPi resistance *via* different mechanisms

To investigate how resistance via loss of PARP1 compares to previously-described PARPi resistance mechanisms in a *BRCA1* mutant context, we used CRISPR-Cas9 mutagenesis to generate SUM149 daughter clones with other previously-identified mechanisms of cellular PARPi resistance, namely reversion mutation in *BRCA1* or deleterious mutations in either *T53BP1* or REV7^22,23,31–33^ (Figure 5a). SUM149 cells with *PARP1* mutations exhibited a comparable if not greater, level of PARPi resistance than these previously identified mechanisms of resistance (Figure 5b, c; p < 0.0001, ANOVA compared to parental cells for all mutants). We compared these models of PARPi resistance to 37 additional breast tumour cell lines using a high throughput drug library screening approach. *PARP1* mutant cells consistently showed the strongest PARPi resistance phenotype across the whole panel, assessed by area under dose-response curve (AUC), while the other resistance mechanisms had slightly lower AUC values similar to breast tumour cell lines without known homologous recombination defects (Figure 5d and Supplementary Figure 4a, b)

**Figure 5.**
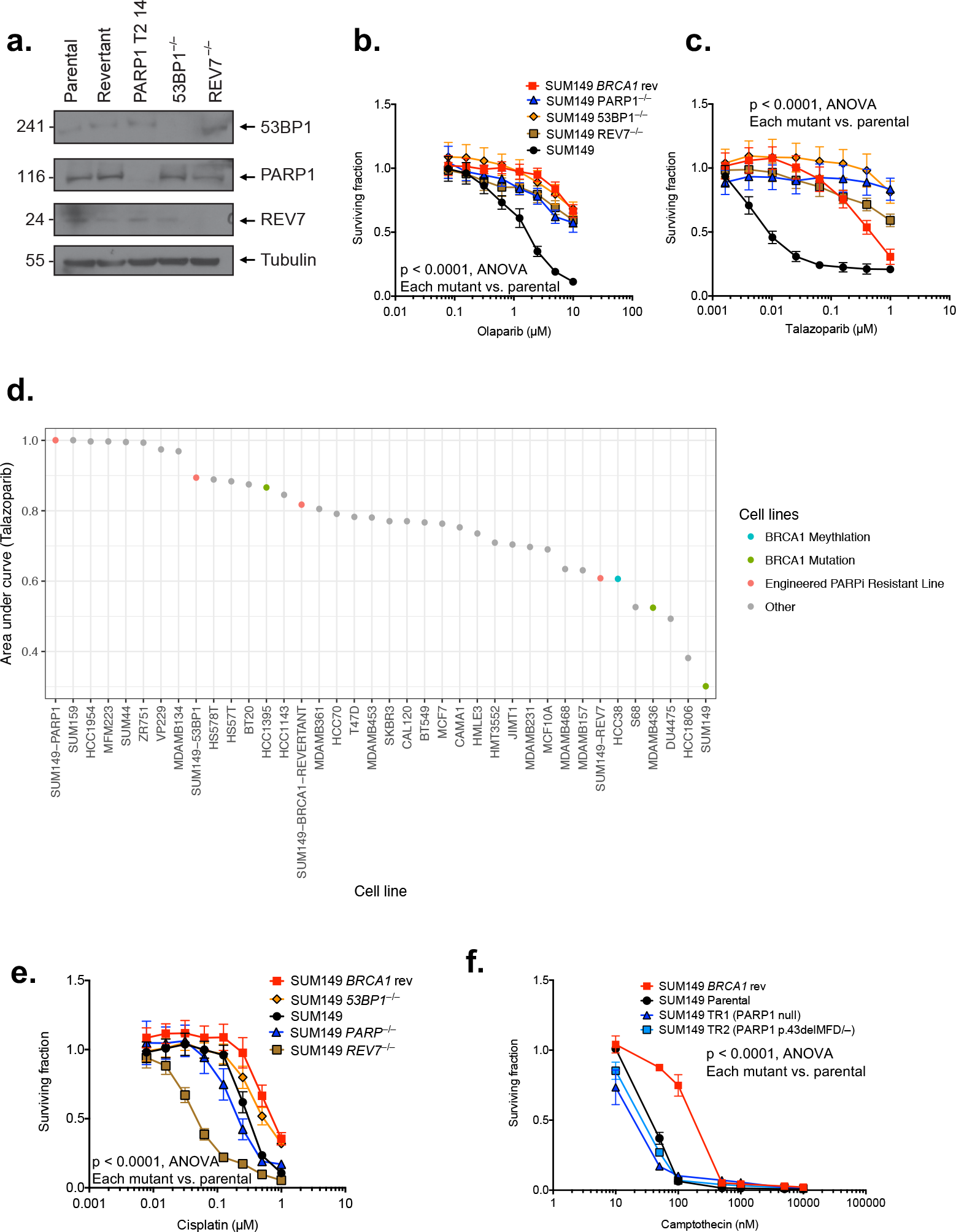
Distinct vulnerabilities of cells acquiring PARPi resistance *via* different mechanisms. **(a)** Western blot showing expression of 53BP1, PARP1 and REV7 protein in a series of isogenic series of SUM149 cell lines generated via CRISPR mutagenesis. Lysates from SUM149 cells (Parental) and clones generated by CRISPR-mediated reversion of the *BRCA1* mutation (“Revertant”), knockout of *PARP1*, *TP53BP1* (53BP1 clones) or *REV7* were blotted and analysed with the indicated antibodies. **(b)** Olaparib and **(c)** talazoparib survival assays for cell lines analysed in (a). Cells were exposed to drug for seven days and survival assayed using CellTiter Glo. All cell lines are significantly resistant to PARPi compared to parental SUM149 cells (p < 0.0001, ANOVA). **(d)** Waterfall plot showing talazoparib area under curve for a panel of breast cancer cell lines exposed to varying concentrations of talazoparib (0.5 – 1000 nM, eight concentrations). **(e)** Differential sensitivity to cisplatin among PARPi resistance mechanisms. *BRCA1* reversion or 53BP1 loss causes resistance to cisplatin relative to the parental cells. However, the *PARP1* mutant clone has similar or slightly increased sensitivity relative to the parental cells and the *REV7* mutant has greatly increased sensitivity (p < 0. 0001, ANOVA, all mutant-parental pairwise comparisons). **(f)** *PARP1* mutant SUM149 cells do not become cross-resistant to the topoisomerase I inhibitor camptothecin, unlike the *BRCA1* revertant clone (p < 0.0001, ANOVA, *BRCA1* revertant compared to parental cells for all drugs).

We assessed whether the particular mechanism of PARPi resistance influenced the sensitivity of these models to additional DNA damaging agents; this could have implications for how patients are treated subsequent to the emergence of PARPi resistance. We found that 53BP1 mutant clones had an intermediate level of resistance to cisplatin (p < 0.0001, ANOVA), as previously described^23^, as did clones with a *BRCA1* reversion, whilst PARP1 mutant cells retained the cisplatin sensitivity of parental SUM149 cells (Figure 5e). In contrast, *REV7* mutants were even more sensitive to cisplatin (p < 0.0001, ANOVA) than all other SUM149 clones, a possible consequence of losing REV7’s role in translesion synthesis^34^ (Figure 5e). Similarly, we noted that whilst a *BRCA1* reversion mutation caused resistance to topoisomerase inhibitors, this was not the case when SUM149 cells possessed PARP1 mutations (Figure 5f, Supplementary Figure 4c, d). This suggested that drug sensitivities in PARPi resistant tumour cells might be predicated in part by the mechanism of PARPi resistance, with different mechanisms of resistance defining secondary vulnerabilities. This implies that understanding the mechanisms of PARPi resistance that emerge in each patient might be important in determining how each individual is subsequently treated.

## Discussion

A key observation that lead to the trapping hypothesis of PARPi cytotoxicity is that, in homologous recombination proficient cells, PARP1 itself is required for cytotoxicty^6,7^. Our genome-wide CRISPR screen revealed that point mutations in the zinc finger domains that abolish DNA binding were sufficient for this resistance, providing direct genetic evidence for the trapping hypothesis.

Our genetic screens also uncovered several clusters of mutations that suggest that regions of PARP1 outside the DNA binding domain can influence trapping (Figures 2d, 4b), observations that are consistent with inter-domain interactions being critical for PARP1 binding and activation^9–11^. These observations are also supported by an orthogonal approach using ethyl methyl sulfonate (EMS) mutagenesis screens in haploid cells (M. Herzog, S.P. Jackson *et al.*, personal communication). These observations add weight to the “reverse allostery” hypothesis proposed by Murai, Pommier *et al,* which suggests that PARPi binding to the catalytic domain of PARP1 allosterically influences interactions between DNA and the N-terminal DNA binding domains of the protein, to the extent that PARP1 becomes trapped on DNA^6^. It is possible that the PARP1 mutations we have identified here are in amino acid residues critical to this reverse allosteric process and therefore prevent allosteric enhancement of DNA binding upon inhibitor binding.

We also isolated a PARP1 mutant, p.848delY, which was associated with PARP inhibitor resistance despite exhibiting some residual PARP1 trapping. This reduced level of trapping might be sufficient to cause resistance – the amount of trapped PARP1 required to induce cytotoxicity is not known. Nor is it known whether all trapping, as observed by chromatin fractionation assays in the presence of alkylating agents, is equal with respect to cytotoxicity – PARP1 trapping at certain sites or lesion types may be more toxic than at others. Alternatively, it is possible that PARP1 trapping is not, in itself, sufficient for cytotoxicity and other factors altered by the mutations are also required. For example, the Y848 residue is part of a solvent exposed helix in PARP1 that may participate in protein-protein interactions – for example with Timeless^35,36^ – that could be important for generating a DNA lesion that is cytotoxic. Mutations in another residue (A925) juxtaposed with Y848 in the PARP1 tertiary structure also led to PARPi resistance, possibly via a similar mechanism (Figure 4e). It is possible that a larger trapped complex of PARP1 and interacting partner(s) is responsible for cytotoxicity, or that trapped PARP1^848delY^ is altered in such a way as to not be cytotoxic. Nevertheless, the resistance phenotype of cells with mutations affecting Y848 at the endogenous *PARP1* locus is slightly less pronounced than a complete PARP1 knockout (Supplementary Figure 2h), supporting the hypothesis that the residual PARP1 trapping corresponds to some residual PARPi cytotoxicity. It will be interesting to study this region further to pinpoint the cause of the reduced cytotoxicity.

Our experiments also showed that *PARP1* mutation can be tolerated in certain *BRCA1* mutant, PARPi sensitive tumour cells. This suggests that PARP1 trapping still underlies the increased cytotoxicity of PARPi in these tumour cells but that some residual BRCA1 function allows these cells to tolerate PARP1 mutations (Figure 3e-g), consistent with previous observations that some *BRCA1* exon 11 mutations do not result in complete loss of BRCA1 function^18,19^. Since *PARP1*;*BRCA1* double mutant cells show distinct drug sensitivities compared to cells that become PARPi resistance via other mechanisms, knowledge of the mechanism of resistance in patients that relapse could inform the best subsequent treatment. Importantly, we also observed a *PARP1* mutation that abolished trapping (R591C, Figure 4f) in a patient with *de novo* resistance to olaparib, suggesting that such mutations can arise in patients and could potentially contribute to resistance. It also seems reasonable to think that PARPi resistance in some patients might display some level of heterogeneity, with multiple different PARPi subclones emerging with distinct mechanisms of resistance; recent advances in the genomic profiling of both solid and liquid biopsies derived from PARPi resistant patients^37–39^ might potentially assess whether this is the case. Whether this turns out to be the case or not, our observation that PARPi resistant cells with different mechanisms of resistance display different chemotherapy sensitivities suggests that defining the molecular features of PARPi resistant disease biopsies might be important to determine the best course of subsequent treatment.

The “tag, mutate and enrich” approach we have used, where the tagging of genes with C-terminal GFP coding sequences (or other selectable genes) followed by the targeted mutagenesis of these genes via CRISPR-Cas9 mediated mutagenesis, allows, in principle, full-length mutants of any gene of interest associated with a selectable phenotype to be identified. This could be employed in the analysis of other resistance mutations observed in patients being treated with targeted therapies in order to annotate likely drivers and passengers of resistance.

## Acknowledgements

We thank Anita Grigoriadis (BCN Research Unit, King’s College, London) for information on *PARP1* copy number in SUM149 cells and Fredrik Wallberg of the ICR, London, for assistance with FACS. This work was funded by Breast Cancer Now and Cancer Research UK Programme Awards to CJL. We acknowledge NHS funding to the NIHR Royal Marsden Hospital Biomedical Research Centre. Funding was also provided by a Stand Up To Cancer— Ovarian Cancer Research Fund Alliance—National Ovarian Cancer Coalition Dream Team Translational Research Grant (grant number SU2C-AACR-DT16-15 to EMS). Stand Up to Cancer is a program of the Entertainment Industry Foundation; research grants are administered by the American Association for Cancer Research, a scientific partner of Stand Up To Cancer.

## Competing interests declaration

CJL and AA are named inventors on patents describing the use of PARP inhibitors and stand to gain from their development as part of the ICR “Rewards to Inventors” scheme.

## Supplementary Tables

**Supplementary Table 1.** CRISPR sgRNA vectors used for the *PARP1* focused mutagenesis screen.

**Supplementary Table 2.** Translated protein alignments for the *PARP1* mutants isolated in the HeLa screen.

**Supplementary Table 3.** Ion Torrent sequencing data for the *PARP1* mutants isolated in the HeLa screen.

**Supplementary Table 4.** Guide sequences for the dense *PARP1* library.

**Supplementary Table 5.** *PARP1* mutations identified in talazoparib-resistant SUM149 *PARP1-tagGFP2* cells in the dense mutagenesis screen.

**Supplementary Table 6.** Primer sequences with Ion Torrent adapters for genotyping mutations in the dense *PARP1* mutagenesis screen.

## Methods

### Cell lines

Mouse ES cells were cultured in Knockout DMEM containing 15% FBS and LIF as previously described^40^. HeLa, CAL51, COV362 and MDA-MB-436 cells were maintained in DMEM supplemented with 10% FBS (with 10 μg/ml insulin in the case of MDA-MB-436). The HeLa cell line expressing PARP1-GFP gene from a bacterial artificial chromosome has been previously described^41^. SUM149PT cells (referred to as SUM149) were maintained in Ham’s F-12 medium supplemented with 5% FBS, 10 μg/ml insulin and 1 μg/ml hydrocortisone.

All cell line identities were confirmed by STR typing and verified free of mycoplasma infection using Lonza MycoAlert.

### Genome-wide mouse CRISPR screen

We screened a previously described library of mouse embryonic stem (ES) cells infected with a lentiviral short guide RNA (sgRNA) library targeting 19,150 mouse genes (average five sgRNAs/gene)^42^. Two million CRISPR mutagenised cells were exposed to a normally lethal concentration (25 nM) of the clinical PARPi talazoparib^15^ (a.k.a. BMN 673; Figure 1a) for six days and allowed to form colonies. Colonies were picked and expanded in 96-well plates. sgRNA sequences from resistant clones were amplified using U6-F and CRISPR-scaf-R primers. PARP1 genotyping of resistant clones coming from the screen was done with Parp1_CRISPRseq-F and Parp1_CRISPRseq-R primers using ReadyMix Taq polymerase (Sigma) and the following conditions: 94 °C for 30 s,30 cycles of 94 °C for 10 s, 56 °C for 10 s and 72 °C for 30 s, and the final extension,72 °C for 4 min.

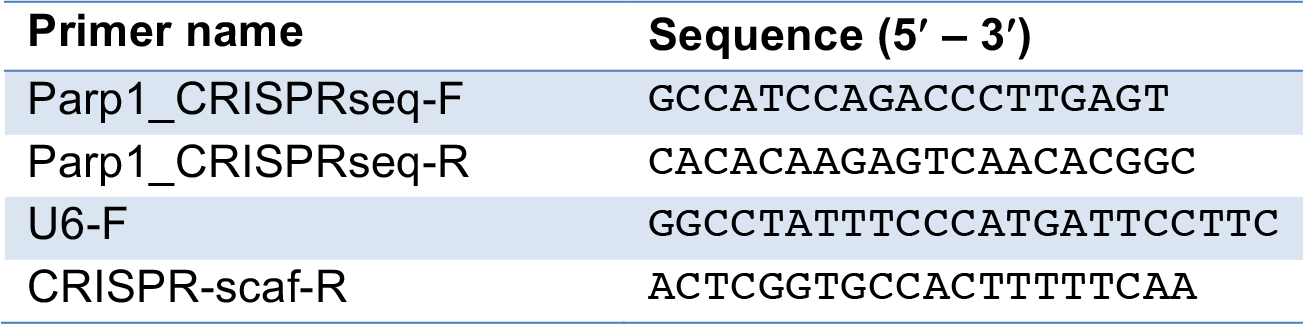

### Genome-wide CRISPR screen in SUM149 cells

SUM149-Cas9 cells were generated by transduction of SUM149 cells with a Cas9-bsd lentivirus and selection in 7 μg/ml blasticidin. Cells were infected at MOI 0.3 with a previously published genome-wide human lentiviral CRISPR library^21^. Cells were selected with puromycin and then placed under talazoparib selection at a concentration that killed all non-infected SUM149 cells (100 nM). Twelve surviving colonies were picked and analysed for the presence of sgRNA sequences by PCR and Sanger sequencing as above.

### Genetically engineered cell lines

CAL51 *PARPT*^−/−^ cells were generated using the Edit-R Gene Engineering System (GE Dharmacon). Cells were seeded at a density of 1 × 10^5^ cells per well in 24-well plates. After 24 hours, cells were transfected with 1 μg Edit-R CRISPR-Cas9 Nuclease Expression Plasmid mixed with 2.5 μl of 20 μM PARP1 T2 crRNA (GAC CAC GAC ACC CAA CCG GAG UUU UAG AGC UAU GCU GUU UUG) and 2.5 μl of tracrRNA (20 μM), using Lipofectamine 3000 according to the manufacturer's instructions (Life Technologies). Three days after transfection, cells were selected in 100 nM talazoparib for five days, and surviving cells FACS sorted into 96-well plates at one cell per well in drug-free medium. After two weeks, the medium was changed to 100 nM talazoparib and cells kept under selection for another month. Targeted genome modifications were analyzed by Sanger sequencing of PCR products cloned into pCR-TOPO-blunt (Life Technologies). Ten individual colonies were sequenced. Clone T2.4, referred to as CAL51 PARP1^−/−^, lacks PARP1 expression by Western blot and has bi-allelic out-of-frame deletions at the target site: a 50 bp deletion and a single base ‘C’ deletion respectively.

Stable Cas9-expressing human cell lines were generated by transducing cells with a Cas9-blasticidin lentivirus^43^ and selecting cells with integrated virus by culturing in medium containing 7 μg/ml blasticidin.

SUM149 TR1 and TR2 were generated by infecting SUM149 Cas9-expressing cells with a pLentiGuide-puro^43^ vector expressing an sgRNA targeting ZnF1 (Hs_PARP1_DBD_cr) and selecting cells with 100 nM talazoparib for 7 days. Single cells were sorted using FACS and the resulting clones genotyped by PCR with Hs_PARP1DBD-genoF and Hs_PARP1DBD-genoR primers using Q5 High-Fidelity DNA Polymerase (NEB) with the following conditions: 98 °C for 30 s, 30 cycles of 98 °C for 10 s, 63 °C for 10 s and 72 °C for 30 s, and the final extension, 72 °C for 2 min.

SUM149 PARP1-tagGFP2 cells were generated by CRISPaint^30^ using as a gene-specific target sgRNA sequence 5’-GCAATTTTAAGACCTCCCTG. GFP-positive single-cell clones were isolated. They were genotyped with the Hs_PARP1_intron22_F and tagGFP2_R primers (98 °C for 30 s, 30 cycles of 98 °C for 10 s, 65 °C for 10 s and 72 °C for 30 s, and the final extension, 72 °C for 2 min), and the PCR product was Sanger sequenced to ensure a single in frame integration. Clones 5 and 8 originated from independent integration events.

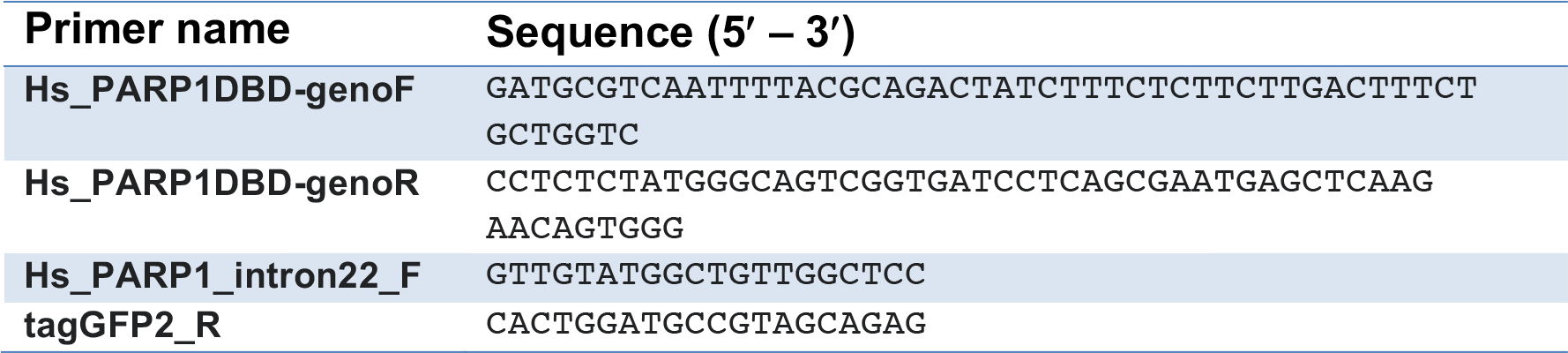
Genotyping primers for SUM149 cells.

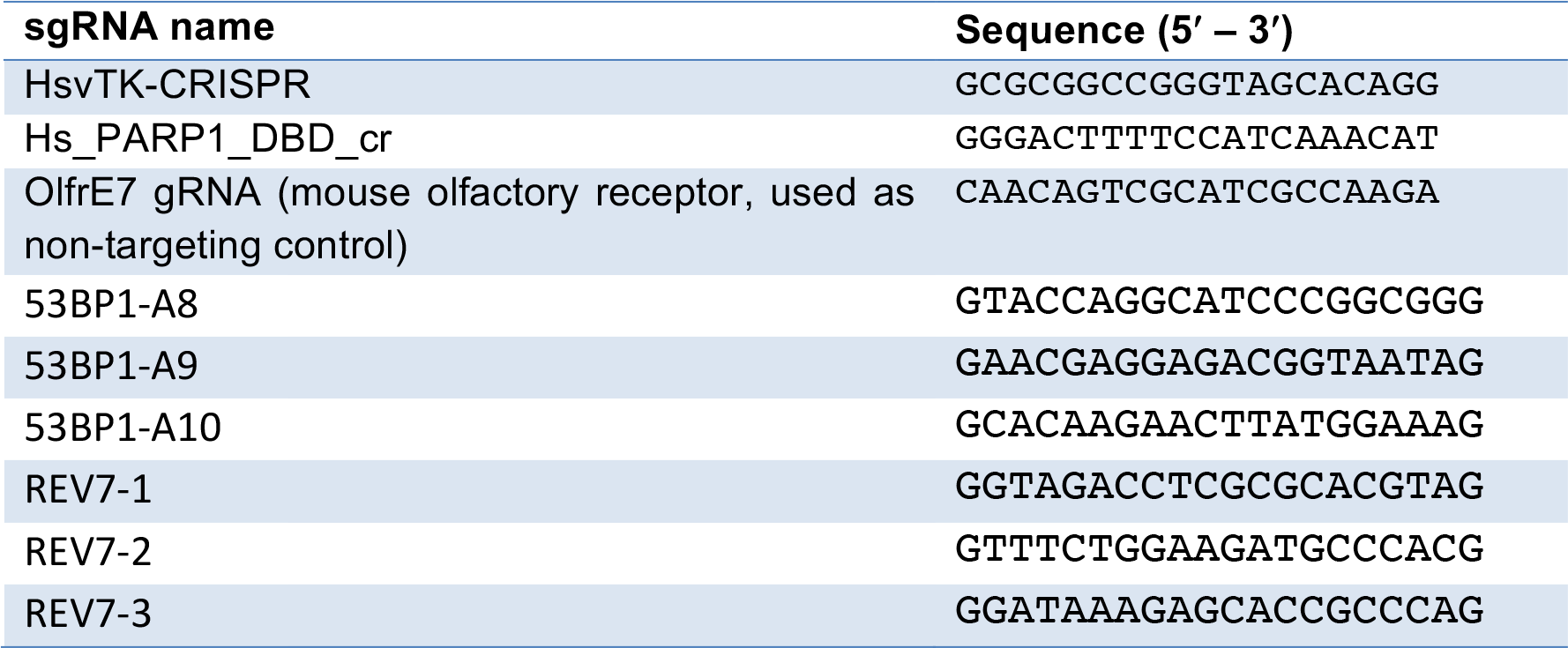
CRISPR guide sequences for individual knockouts and controls.

### Reagents

Drugs, including olaparib and talazoparib, were purchased from Selleckchem, except temozolomide and methyl methanesulfonate (Sigma-Aldrich). Antibodies for PARP1 (Cell Signaling #9542), PAR (Trevigen, 4335-AMC-050 and Enzo 10H), GFP (Sigma-Aldrich, 11814460001), Top1 (BD #556597), BRCA1 (IP: Santa Cruz sc6954; Blot: MS110, Abcam ab16780), Tubulin (Sigma, T6074), REV7 (MAD2L2, Abcam, ab180579).

### Cell viability and clonogenic assays

The viability of cells was measured after five days exposure to various concentrations of drugs using the Cell Titre-Glo assay (Promega). Long-term drug exposure effects were assed by colony formation assay after 7-10 days exposure to a drug as described previously^1^, with the modification that cells were stained at the end of the assay with sulforhodamine B (SRB). When plotting survival curves, the surviving fraction was calculated relative to DMSO (solvent) exposed cells.

### PARP1 sgRNA screen in HeLa cells

29 sgRNAs were designed using the ChopChop algorithm^44^. The guides were synthesized and cloned in an array format into pLentiGuide-puro^43^ by Eurofins genomics. A single colony for each vector was used to inoculate a pooled culture for plasmid maxi prep (Qiagen). Six such pools of vectors were packaged into viruses as previously described^45^. These pooled viruses were used to infect HeLa PARP1-GFP/Cas9 cells. sgRNA expressing cells were enriched with a three day puromycin selection (3 μg/ml). The cells were subsequently incubated in 1 μM talazoparib for 12 additional days. The GFP-positive population was enriched by cell sorting (BD FACSAria). Total RNA was extracted from each of the six pools. One microgram of total RNA was converted to cDNA with SuperScript III (ThermoFisher) and a GFP-specific primer (5’- CCT TGA TGC CGT TCT TCT GCT TG), which allowed for amplification only of the target *PARP1-GFP* transcripts. Each of the six pooled cDNAs was amplified with its corresponding Ion Torrent adapted primer pair (Figure 2a) flanking the gRNA target sites for that pool. This amplification used Q5 High-Fidelity DNA Polymerase (NEB) with the following PCR conditions: 98 °C for 30 s, 30 cycles of 98 °C for 10 s, 64 °C for 10 s and 72 °C for 30 s, and the final extension, 72 °C for 2 min. Primer sequences are provided in the table below, where the Ion Torrent sequencing adapters are represented by A1 (CCA TCT CAT CCC TGC GTG TCT CCG A) and P1(CCT CTC TAT GGG CAG TCG GTG ATC).

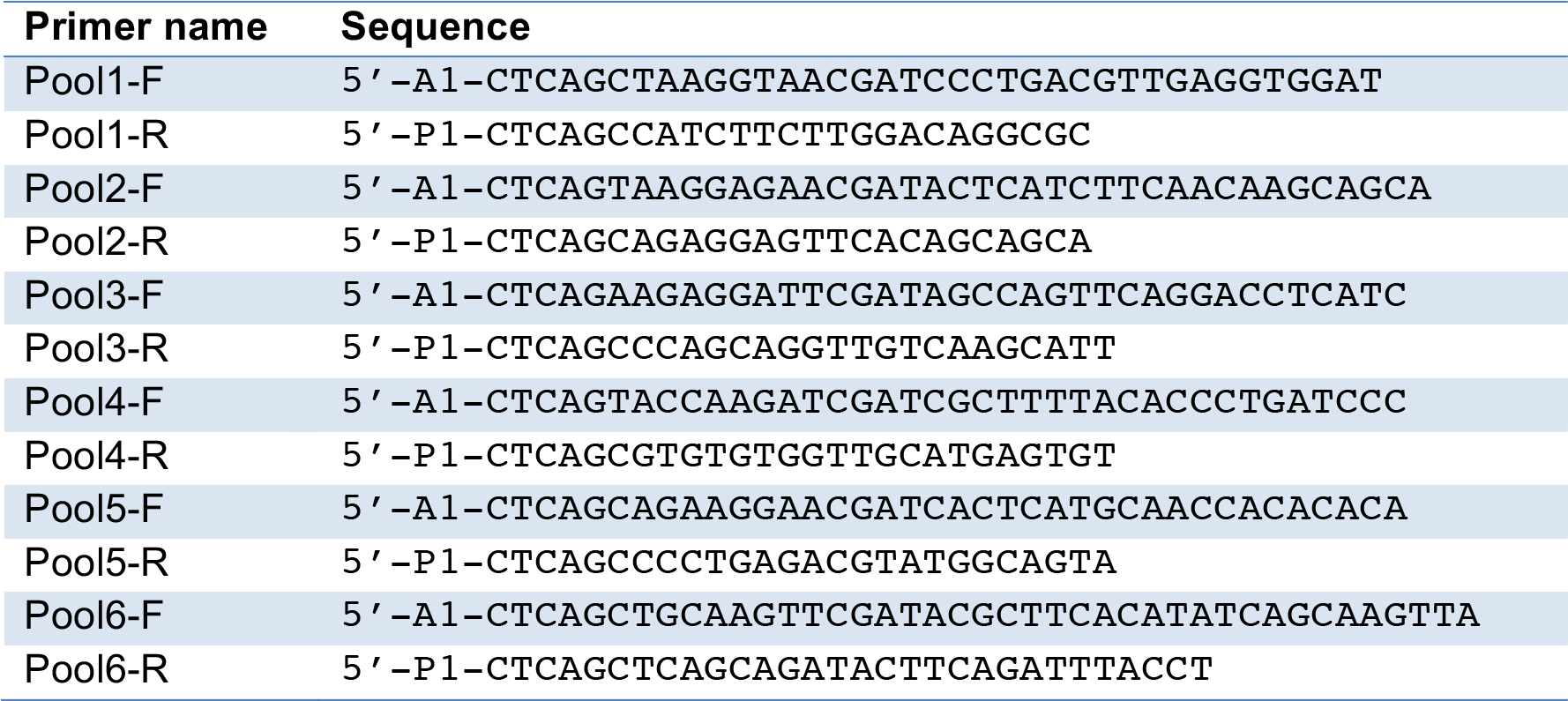

### PARP1 dense screen in SUM149 cells

A dense CRISPR library comprising 489 guides targeting *PARP1* was synthesised (Twist biosciences, Supplementary Table 4) and cloned into the BbsI site of pKLV5-U6gRNA5-PGKPuroBFP^21^. One million cells from each of two SUM149 PARP1-tagGFP2 clones were infected at MOI 0.3 as above and selected in puromycin. Cas9 was induced by treatment with doxycycline for two days after which cells were replated and 100 nM talazoparib added. cDNA was prepared using a GFP primer as above and fifteen overlapping amplicons prepared by PCR with Ion Torrent adapter tailed primers for genotyping (Supplementary Table 5).

### Sequencing and analysis

Purified RT-PCR products were mixed and sequenced using the Ion Torrent PGM and a 318 chip with 850 flows. Data were converted to FASTQ format and aligned to the *PARP1* cDNA sequence (ENST00000366794) using Novoalign (Novocraft technologies). Mutations were called from the alignments using the Ensembl Variant Effect Predictor REST API^46^ (implementation:github.com/GeneFunctionTeam/bioruby-sam-mutation). Sequences were also translated and multiple alignments generated using Clustal Omega (http://www.ebi.ac.uk/Tools/msa/clustalo/). For the dense screen, coverage was calculated per base using samtools pileup (github.com/samtools).

### Microirradiation assays

Cells were grown in glass-bottom culture dishes (MaTek, P35G-0.170-14-C) in 10% FBS DMEM media and maintained at 37°C and 5% CO_2_ in an incubation chamber mounted on the microscope. Imaging was carried out on Andor Revolution system, 60x water objective with micropoint at 365 nm. Only cells with similar GFP signal intensity were measured. The background intensity (in the vicinity of the microirradiation area in the nucleus) was subtracted from that at the microirradiation point and the maximum was normalised to 1.

### Chromatin fractionation (trapping assay)

The chromatin fractionation assay for PARP trapping was based on a previously published protocol^6^. Cells were grown in 6-well plates, exposed for 1-4 hours to 500 nM talazoparib and 0.01% MMS and fractionated with the Subcellular Protein Fractionation kit for Cultured Cells (ThermoFisher #78840) according to the manufacturer’s recommendations. Equivalent volumes of each fraction were analysed by western blot using the PARP1 (Cell Signaling #9542, rabbit), with TOP1 (BD #556597, mouse) or Histone H3 (CST #9717, rabbit; low molecular weight region of blot probed separately) as fractionation controls. Blots were imaged using the LiCor Odyssey multicolour imaging system and secondary antibodies (Donkey anti-rabbit 680 and donkey antimouse 800).

### Protein purification

PARP1 point mutations were introduced in an IPTG-inducible C1-PARP1 plasmid (expressing His-MBP tagged human PARP1 cDNA) and transformed in Rosetta *E. coli* (Novagen). Cells were grown in terrific broth at 37°C until reaching OD_600_ 1.0, 100 mM ZnSO_4_ was added and cells further grown to OD_600_ 2.0. 0.5 mM IPTG was added and cells were shaken at 18°C over night. Pellets were collected by centrifugation and stored at −80°C. Pellets were lysed (50 mM Tris-HCl pH 7.5, 500 mM NaCl, 10 mM imidazole, 10 mM mercaptoethanol, protease inhibitors), sonicated and cleared by centrifugation and 0.45 μm filtration. The lysate was passed through a 5 ml His Trap column (GE). The column was washed and bound proteins eluted with a linear imidazole gradient. Pooled PARP1-containing fractions were dialyzed over night (50 mM Tris-HCl pH 7.5, 250 mM NaCl, 5 mM mercaptoethanol) in the presence of MBP-TEV protease. Dialyzed proteins were cleared through 0.45 μm filter and cleaved MBP and MBP-TEV were removed through another 5 ml His Trap column. Samples were loaded on a 5 ml HiTrap Heparin column (GE) and eluted with a linear salt gradient. PARP1-containing fractions were pooled, concentrated to 1 ml and injected on a HiLoad 16/600 Superdex 200 coulmn (GE) equilibrated in 25 mM HEPES-NaOH pH7.5, 100 mM NaCl, 2 mM TCEP. PARP1-containing fractions were pooled, concentrated and snap-frozen in liquid N_2_.

### PARylation assay

DNA dumbbell ligand (DB4) was prepared as previously described^10^. *In vitro* PARylation reactions were carried out in a buffer (50 mM HEPES pH 7.5, 150 mM NaCl, 10 mM MgCl_2_ and 1 mM DTT) containing 100 nM PARP1 protein and 1 μM DB4. The reaction was started by the addition of NAD^+^ to a 225 μM final concentration (spiked with 5 μCi/ml ^32^P-NAD^+^) and incubation at 30°C. Aliquots were taken at various time points and the reaction was stopped by the addition of 2x SDS-PAGE loading dye and heat denaturation. Samples were resolved on SDS-PAGE, dried and exposed on a phosphoimager plate (GE). Total radioactive intensity was quantified in the lanes between the well start and the position of PARP1.

### siRNA transfection

Lipofectamine RNAimax was used according to the manufacturer’s instructions (Invitrogen). Cells were transfected with Smart Pools targeting the appropriate gene (GE Dharmacon).

### Patient tumour sequencing

The *PARP1*:R591C mutation was identified in the archival tumour sample of a 69 year old patient (without a *BRCA1* or *BRCA2* germline mutation) with platinum-resistant high grade serous ovarian cancer enrolled in a phase I trial of olaparib and durvalumab (NCT02484404). A panel of cancer susceptibility and DNA repair genes (BROCA-HR^47^) was sequenced and the *PARP1*.R591C mutation (genomic: chr1:226,564,979G>A) was identified at a variant allele frequency (VAF) of 0.33 (65/198 reads). This ovarian carcinoma also had a TP53 mutation (p.R196*) with VAF of 0.72. The patient did not have a response to olaparib and durvalumab based on RECIST v1.1 criteria.

